# iPSC-derived hepatocytes from patients with nonalcoholic fatty liver disease display a disease-specific gene expression profile

**DOI:** 10.1101/2020.04.20.052001

**Authors:** Caroline C. Duwaerts, Chris L. Her, Nathaniel J. Phillips, Holger Willenbring, Aras N. Mattis, Jacquelyn J. Maher

**Affiliations:** Departments of Medicine, University of California San Francisco, San Francisco, CA, USA, 94143; Departments of Surgery, University of California San Francisco, San Francisco, CA, USA, 94143; Departments of Pathology, University of California San Francisco, San Francisco, CA, USA, 94143; Departments of Liver Center, University of California San Francisco, San Francisco, CA, USA, 94143; Technology Center for Genomics & Bioinformatics, University of California Los Angeles, Los Angeles, CA, USA, 90095

**Keywords:** steatohepatitis, insulin resistance, lipid, alpha-arrestin, thioredoxin

## Abstract

Nonalcoholic fatty liver disease (NAFLD) is one of the leading causes of liver disease worldwide.1 Animal models are widely used to investigate the mechanisms of fatty liver disease, but they do not faithfully represent NAFLD in humans.2 Thus, there is strong interest in studying NAFLD pathogenesis directly in humans whenever possible. One strategy that is gaining momentum is to utilize iPSC-derived hepatocytes from individual human subjects in complex cell/organ platforms with the goal of reproducing a NAFLD-like state in vitro.3-6 Our group has taken a different approach, positing that iPSC-Heps from a population of NAFLD patients would provide independent insight into the human disease. In this study we generated iPSCs and iPSC-Heps from a well-defined cohort of NAFLD patients. Our objective was to determine whether as a group, in the absence of any metabolic challenge, they exhibit common disease-specific signatures that are distinct from healthy controls.

## INTRODUCTION

Nonalcoholic fatty liver disease (NAFLD) is one of the leading causes of liver disease worldwide.^1^ Animal models are widely used to investigate the mechanisms of fatty liver disease, but they do not faithfully represent NAFLD in humans.^2^ Thus, there is strong interest in studying NAFLD pathogenesis directly in humans whenever possible. One strategy that is gaining momentum is to utilize iPSC-derived hepatocytes from individual human subjects in complex cell/organ platforms with the goal of reproducing a NAFLD-like state in vitro.^3-6^ Our group has taken a different approach, positing that iPSC-Heps from a population of NAFLD patients would provide independent insight into the human disease. In this study we generated iPSCs and iPSC-Heps from a well-defined cohort of NAFLD patients. Our objective was to determine whether as a group, in the absence of any metabolic challenge, they exhibit common disease-specific signatures that are distinct from healthy controls.

## METHODS

Please see Supplementary Material.

## RESULTS

iPSCs were generated from peripheral blood mononuclear cells collected from 13 unrelated patients with biopsy-proven NAFLD and 14 healthy control subjects. Individual clinical data for all 27 subjects are reported in **Table S1**. Two of the 13 NAFLD patients had isolated hepatic steatosis and 11 had histologic NASH. Of the 11 patients with NASH, 9 (82%) had advanced fibrosis. Seven NAFLD patients (54%) and 3 controls (21%) were homozygous for the *PNPLA3* G allele (*P* = 0.08).

iPSC-Heps differentiated from control and NAFLD iPSCs exhibited similar morphology, including lipid content (**Figure S1A**). All iPSC-Heps displayed high-level expression of hepatocyte-specific genes following differentiation (**Figure S1B**). Transcriptomic analysis of iPSCs before differentiation revealed no clustering by disease status (**Figure 1A**); differentiated iPSC-Heps, however, segregated into two groups based on the presence or absence of NAFLD. The principal disease and function pathways up-regulated in NAFLD iPSC-Heps highlighted inflammation and stress responses (**Figure 1B**). Similarly, upstream regulators induced in NAFLD iPSC-Heps featured several genes controlling pro-inflammatory pathways (**Figure S1C**).

**Figure 1.**
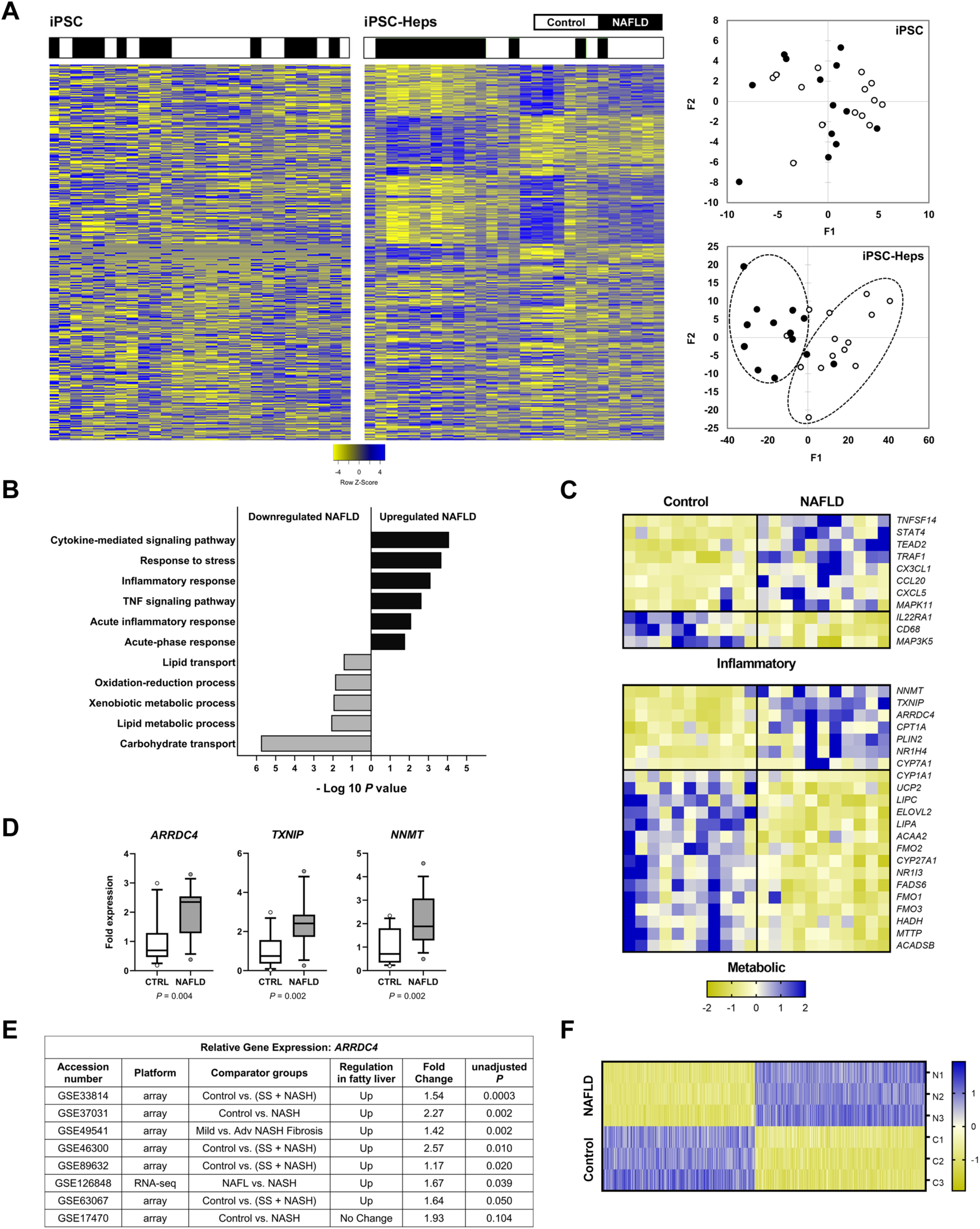
Comparison of the transcriptomic profiles of iPSC and iPSC-Heps from NAFLD and control subjects. (A) Heatmaps illustrate unsupervised clustering of RNA sequencing data from control and NAFLD iPSCs (left panel) and control and NAFLD iPSC-Heps (right panel). The heatmaps, as well as the accompanying principal component analyses, show clustering of iPSC-Heps but not iPSCs by disease group. (B) Graph illustrates select Gene Ontology pathways that were significantly up-or down-regulated in NAFLD vs. control iPSC-Heps. (C) Heatmaps show differential regulation of genes in two major categories within the top 500: inflammatory molecules and proteins involved in metabolism of nutrients and xenobiotics. (D) The top three metabolic genes up-regulated in the metabolism category (fold change > 2.5, *P* < 0.0005) were confirmed by QPCR. (E) *ARRDC4* was identified as being significantly up-regulated in NAFLD vs. control subjects from 7 of 8 cohorts registered in the Gene Expression Omnibus (GEO) repository. (F) Heatmap illustrates the reproducibility of gene expression across several independent iPSC-Hep differentiations from one NAFLD and one control iPSC line. 5,755 differentially expressed genes are represented.

Among the top 500 genes differentially expressed between NAFLD and control iPSC-Heps (*P* < 0.003) were candidates related to injury and inflammation as well as metabolism (**Figure 1D**). Genes up-regulated in NAFLD iPSC-Heps included select chemokines and pro-inflammatory signaling molecules (*CX3CL1, CCL20, CXCL5, STAT4, TRAF1, MAPK11*); the IL22 receptor (*IL22RA1)*, which promotes survival signaling, was down-regulated. Unexpectedly, ASK1 (*MAP3K5*), a stress kinase whose activity has been implicated as a driver of experimental NAFLD, was down-regulated in NAFLD iPSC-Heps.

In the category of metabolism, iPSC-Heps exhibited up-regulation of two related genes: arrestin domain-containing 4 (*ARRDC4*) and thioredoxin inhibitory protein (*TXNIP*). ARRDC4 and TXNIP belong to a family of proteins termed alpha-arrestins, whose expression increases during insulin resistance.^7^ Another gene up-regulated in NAFLD iPSC-Heps encodes nicotinamide N-methyltransferase (*NNMT*): NNMT regulates cellular levels of S-adenosylmethionine and NAD+, and its expression correlates positively with obesity and fatty liver. Also notable in the metabolism category was that several genes involved in lipid disposal were down-regulated in NAFLD iPSC-Heps. Some of the downregulated genes promote fatty acid beta-oxidation (*ACAA2, HADH, ACADSB*), others generate polyunsaturated fatty acids (*ELOVL2, FADS6*), and still others promote lipid export from hepatocytes (*MTTP)*.

The top three metabolic genes up-regulated in NAFLD iPSC-Heps by RNA-seq (fold change > 2.5, *P* < 0.0005) were confirmed by QPCR (**Figure 1D**). The GEO Profiles database was queried to determine whether these genes were similarly up-regulated in other fatty liver disease cohorts. *ARRDC4* was significantly up-regulated in 7 of 8 queried populations (**Figure 1E**). *TXNIP* and *NNMT* were not consistently up- or down-regulated (not shown).

To determine the reproducibility of the gene expression profile from our subjects across several iPSC differentiations, we performed RNA sequencing on three independently differentiated iPSC-Hep cultures from a NAFLD patient and three from a control subject, each of which originated from a single iPSC line. Differentially expressed genes between the NAFLD and control iPSC-Heps revealed excellent reproducibility across separate differentiations, as well as the persistence of a distinction between NAFLD and control iPSC-Heps over time (**Figure 1F**).

## DISCUSSION

Our results show that iPSC-Heps from NAFLD patients display a gene signature that correlates with the presence of liver disease in the donor. This disease-specific signature was not present in undifferentiated iPSCs, but appeared only upon differentiation to hepatocyte-like cells. The fact that a NAFLD phenotype became discernable in iPSC-Heps even in the absence of a metabolic challenge indicates that NAFLD patients harbor a predilection to disease that manifests specifically within hepatocytes.

At the pathway level, the NAFLD signature in iPSC-Heps was dominated by genes associated with stress responses and inflammation. This underscores that NAFLD is a disease marked by cellular dysfunction and repair. Metabolic alterations were not prominent in NAFLD iPSC-Heps at the pathway level, but they emerged upon scrutiny of individual differentially expressed genes. In this fashion we identified enhanced expression of genes that portend insulin resistance and mitochondrial dysfunction, as well as reduced expression of genes involved in lipid oxidation and lipid secretion.

An intriguing finding was that two of the most significantly up-regulated genes in NAFLD iPSC-Heps, *ARRDC4* and *TXNIP*, are members of the alpha-arrestin family and that *ARRDC4* is also up-regulated in other NAFLD cohorts. *TXNIP* is already known for its ability to promote oxidant stress and cell death, and thus it may play a dual role in NAFLD pathogenesis. Further exploration of these and other differentially expressed genes is underway.

Our experiments are distinct from, but complementary to the work of others who are using iPSCs from individual subjects to investigate the pathogenesis of metabolic liver diseases.^3-6, 8^ An important benefit of our cohort approach is its potential to identify novel characteristics that are diagnostic or even predictive of NAFLD in a larger population. The fact that we uncovered NAFLD clustering even in a small cohort is encouraging.

In summary, the current experiments establish that iPSC-Heps are a tractable model for identifying hepatocyte-specific attributes that contribute to human NAFLD. The data indicate that iPSC-Heps from unrelated NAFLD patients share characteristics that portend a disease phenotype. The reproducibility of the NAFLD signature bodes well for expanded study of iPSC from larger, more diverse NAFLD populations. Further development of this cohort-based iPSC approach stands to provide novel insight into human NAFLD.

## FUNDING

This work was funded by research grants from the California Institute for Regenerative Medicine (IT1-06563), AbbVie, Inc. and NIH: R21 DK118380 (JJM), UG3 DK120004 (HW) and K08 DK098270 (ANM). Additional support was provided by core facilities within the UCSF Liver Center (P30 DK026743).

## CONFLICT OF INTEREST STATEMENT

The authors declare no competing interests.

## AUTHOR CONTRIBUTIONS

Conceptualization: JJM

Methodology: CCD, ANM, JJM, HW

Investigation: CCD, CLH, JJM, NJP

Formal analysis: CCD, JJM, NJP

Resources: ANM, JJM, HW

Writing – original draft: CCD, JJM

Writing – review and editing: CCD, ANM, JJM, NJP, HW

Visualization: CCD, JJM, NJP

Supervision: JJM

Funding acquisition: JJM

## ACKNOWLEDGMENTS

The authors are indebted to Eric Hoffman and Kevin Siao, who provided valuable assistance with subject recruitment and experimentation, and to Dr. Stephen A. Duncan, who provided advice and assistance with iPSC-Hep differentiation.

## Supplementary Material

## METHODS

### Subject recruitment

NAFLD patients were recruited from the Liver Clinic at the Zuckerberg San Francisco General Hospital and Trauma Center (ZSFG). Inclusion criteria were age ≥ 18 years, alcohol consumption < 1 drink per day for women and 2 drinks per day for men, a confirmed diagnosis of NAFLD by liver biopsy, and the ability to provide informed consent. Exclusion criteria were the presence of other liver diseases (hepatitis B, hepatitis C, autoimmune or hereditary liver disease) or the presence of active infection with human immunodeficiency virus (HIV). Control subjects were recruited by local advertisement. Inclusion criteria were age ≥ 18 years, BMI < 30, alcohol consumption < 1 drink per day for women and 2 drinks per day for men, the absence of known liver disease or infection with hepatitis B virus (HBV), hepatitis C virus (HCV) or HIV, and the ability to provide informed consent. Control subjects were pre-screened for HBV, HCV and HIV and for liver disease based on serum levels of aspartate aminotransferase, alanine aminotransferase and total IgG. Only those with normal results for all tests were permitted to serve as controls. Controls were 50% male and NAFLD patients were 38.5% male (*P* = 0.57). Mean BMI for controls was 24.8 (19.0-29.8), and mean BMI for NAFLD patients was 32.7 (22.1-55.9) (*P* = 0.003). All enrollees (control and NAFLD) submitted demographic and health-related information via written questionnaire. In addition, for the NAFLD patients, clinical and histologic data relevant to NAFLD were obtained from the medical record at ZSFG. All subjects underwent blood collection for SNP genotyping and iPSC reprogramming. The study was approved by the Institutional Review Board as well as the Human Gamete, Embryo and Stem Cell Research Committee at the University of California San Francisco.

### Generation of iPSC and iPSC-Heps

Peripheral blood mononuclear cells were used to generate iPSC from all subjects. iPSC were generated by Fujifilm Cellular Dynamics International, Inc. (Novato, CA) using a virus-free, episomal reprogramming protocol as described by Yu and colleagues (Yu et al., 2011). iPSC passed quality control testing for pluripotency and loss of reprogramming factors. iPSC-Heps were differentiated from iPSC using a minor modification of the protocol developed by Dr. S. A. Duncan (Si-Tayeb et al., 2010). Precursor iPSC were harvested for RNA extraction and RNA sequencing immediately preceding hepatocyte differentiation. The corresponding iPSC-Heps were harvested on day 22 of the differentiation process, at which time they exhibited characteristic features of hepatocyte-like cells. iPSC-Heps were not exposed to any metabolic or toxic stimuli prior to harvest. RNA samples were submitted to the Technology Center for Genomics & Bioinformatics at UCLA for library construction and RNA sequencing using the Illumina HiSeq 3000 Sequencing System. 30 million reads were obtained per sample, 60-70% of which mapped to the transcriptome Transcriptomic data were analyzed on UCLA’s Hoffman2 distributed computing cluster. SNP genotyping and QPCR were performed using TaqMan assays (Thermo Fisher).

### Quantification and Statistical Analysis

Reads were mapped to UCSC’s hg19 transcript set using Bowtie 2 version 2.1.0 (Langmead and Salzberg, 2012). Gene expression was quantified using RSEM’s (v1.2.15) expectation maximization algorithm (Li and Dewey, 2011). Quantified gene counts were first normalized to counts per million. The counts per million were then further normalized using the Trimmed mean of M-values (TMM) functionality in edgeR (Robinson et al., 2010). Differential gene expression between conditions was calculated by fitting a generalized negative binomial log-linear model using edgeR (Robinson et al., 2010). Genes that were considered differentially expressed contained a cpm count greater than 1 across at least half the replicates in the given comparison in addition to the standard *P* < 0.05, |FC| > 1.5 constraints. Differentially expressed genes were given to IPA (Kramer et al., 2014), which created lists of Canonical Pathways, Upstream Regulators, Diseases and Functions, and Networks. These lists were based on the fold change of the differentially expressed genes. Additional comparisons were made between iPSC and iPSC-Heps and between control and NAFLD iPSC-Heps using open-source software from the database for annotation, visualization and integrated discovery (DAVID) and the Search Tool for the Retrieval of Interacting Genes/Proteins (STRING).

Custom heatmaps of the RNAseq data were created by calculating the coefficient of variation https://en.wikipedia.org/wiki/Coefficient_of_variation across all samples, and then taking the top “N” genes with the largest coefficient of variation. Unsupervised, hierarchical clustering was performed at the gene level by using the single-linkage algorithm between samples’ TMM-normalized counts per million. Additional heatmaps highlighting specific gene subsets were generated using Morpheus open software from the Broad Institute https://software.broadinstitute.org/morpheus/. Samples in the latter heatmaps were ordered based on Pearson’s correlation coefficient.

### Data availability

RNA-seq data deposited in GEO: GSE138312.

---

**Table S1.** Clinical description of each participant.

| Age | Sex | BMI | DM2 | PNPLA3<br>Genotype | Steat | Bal | Inf | NAS | Fib | Pathologist<br>Diagnosis |
| --- | --- | --- | --- | --- | --- | --- | --- | --- | --- | --- |
| **Control** |  |  |  |  |  |  |  |  |  |  |
| 23 | F | 18.1 | N | GC | -- | -- | -- | -- | -- | -- |
| 23 | F | 21.1 | N | GG | -- | -- | -- | -- | -- | -- |
| 31 | M | 29.8 | N | CC | -- | -- | -- | -- | -- | -- |
| 32 | F | 25.4 | N | CC | -- | -- | -- | -- | -- | -- |
| 58 | F | 25.6 | N | GC | -- | -- | -- | -- | -- | -- |
| 42 | M | 26.0 | N | GC | -- | -- | -- | -- | -- | -- |
| 38 | M | 21.1 | N | CC | -- | -- | -- | -- | -- | -- |
| 47 | M | 23.1 | N | GC | -- | -- | -- | -- | -- | -- |
| 40 | F | 27.6 | N | CC | -- | -- | -- | -- | -- | -- |
| 38 | M | 26.8 | N | CC | -- | -- | -- | -- | -- | -- |
| 40 | M | 22.1 | N | GC | -- | -- | -- | -- | -- | -- |
| 29 | F | 26.2 | N | GC | -- | -- | -- | -- | -- | -- |
| 31 | F | 27.4 | N | GG | -- | -- | -- | -- | -- | -- |
| 32 | M | 22.4 | N | GG | -- | -- | -- | -- | -- | -- |
| **NAFLD** |  |  |  |  |  |  |  |  |  |  |
| 43 | F | 31.5 | N | GG | 1 | 2 | 1 | 4 | 1 | NASH |
| 56 | F | 33.1 | N | GC | 1 | 2 | 0 | 3 | 4 | NASH |
| 40 | M | 26.9 | N | GG | 3 | 0 | 1 | 4 | 0 | NAFL |
| 58 | F | 55.9 | Y | GG | 3 | 0 | 3 | 6 | 4 | NASH |
| 48 | M | 22.1 | Y | GG | 2 | 1 | 1 | 4 | 3 | NASH |
| 48 | M | 29.8 | N | GC | 2 | 1 | 1 | 4 | 3 | NASH |
| 47 | F | 39.3 | Y | GG | 1 | 2 | 1 | 4 | 3 | NASH |
| 38 | M | 23.4 | N | GG | 1 | 1 | 1 | 3 | 1 | NASH |
| 41 | F | 38.0 | N | GC | 2 | 2 | 1 | 5 | 3 | NASH |
| 45 | F | 26.1 | N | GG | 2 | 0 | 0 | 2 | 0 | NAFL |
| 47 | F | 37.7 | Y | GC | 3 | 1 | 2 | 6 | 3 | NASH |
| 68 | F | 25.8 | Y | GG | 2 | 2 | 2 | 6 | 3 | NASH |
| 52 | M | 31.6 | Y | GG | 2 | 2 | 1 | 5 | 4 | NASH |

Bal, ballooning; DM2, type 2 diabetes; Fib, fibrosis; Inf, inflammation; NAS, NAFLD Activity Score; Steat, steatosis; NAFL, nonalcoholic fatty liver; NASH, nonalcoholic steatohepatitis

**Figure S1.**
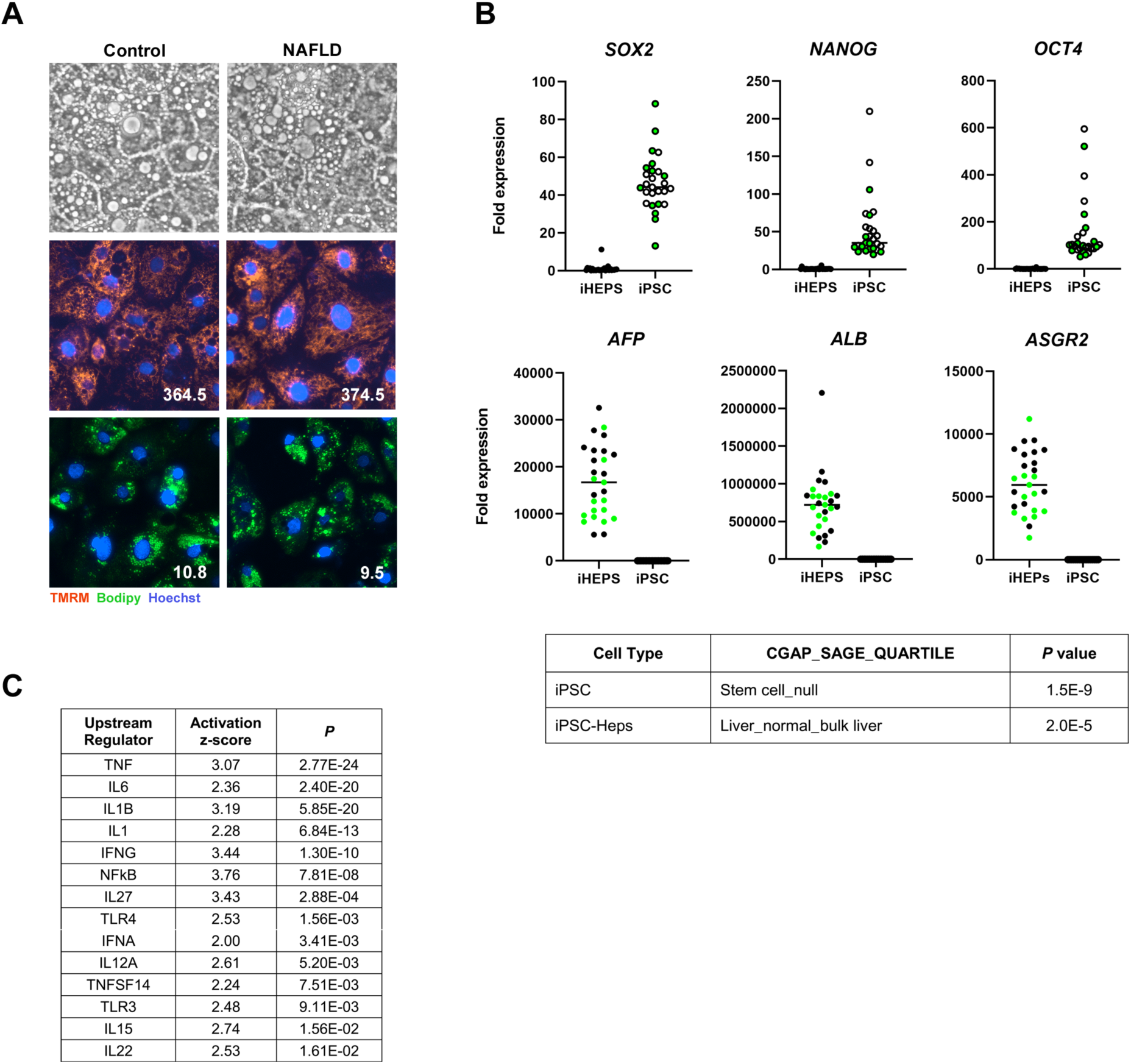
Morphology of iPSC-Heps and confirmation of hepatocyte-specific gene expression following differentiation; differential gene expression of upstream regulatory genes in iPSC-Heps. (A) Photomicrographs illustrate control and NAFLD iPSC-Heps on day 22 of differentiation. Panels depict phase contrast morphology and fluorescent images documenting mitochondrial polarization (TMRM, tetramethylrhodamine methyl ester) and lipid content (Bodipy 493/503). Numbers indicate mean fluorescence intensity per cell nucleus. (B) <u>Top panel</u>: Graphs depict the relative expression of 3 pluripotency genes (*SOX2, NANOG, OCT4*) and 3 liver-specific genes (*AFP, ALB, ASGR2*) measured in iPSC and iPSC-Heps by QPCR. Colored symbols = NAFLD subjects; black symbols = controls. <u>Bottom panel</u>: Cancer Genome Anatomy Project_Serial Analysis of Gene Expression (CGAP_SAGE) analysis of the top 100 differentially expressed genes in iPSC and iPSC-Heps identifies the cells as stem cells and liver, respectively. (C) Table shows upstream regulatory genes identified by IPA as significantly up-regulated in NAFLD vs. control iPSC-Heps.

## Notes

https://www.ncbi.nlm.nih.gov/geo/query/acc.cgi?acc=GSE138312

